# Life on the road: fish communities composition in roadside ditches of the Atlantic Forest

**DOI:** 10.1101/2025.06.18.660392

**Authors:** João Henrique Alliprandini da Costa, Amanda Selinger, Francisco Langeani, Ursulla Pereira Souza, Rafael Mendonça Duarte

**Affiliations:** Laboratório de Ecofisiologia e Toxicologia Aquática, Universidade Estadual Paulista “Júlio de Mesquita Filho” - UNESP, Instituto de Biociências, Praça Infante Dom Henrique, s/n, Parque Bitaru, 11330-900 São Vicente, SP, Brazil., (RMD); Laboratório de Biologia de Organismos Marinhos e Costeiros, Universidade Santa Cecília, Rua Oswaldo Cruz, 277, Boqueirão, 11045-907 Santos, SP, Brazil. (AS), (UPS); Laboratório de Ictiologia, Universidade Estadual Paulista “Júlio de Mesquita Filho” – UNESP, Instituto de Biociências, Letras e Ciências Exatas, Rua Cristóvão Colombo, 2265, Jardim Nazareth, 15054-000 São José do Rio Preto, SP, Brazil.

**Keywords:** Habitat fragmentation, Freshwater fish, Ephemeral habitats, Artificial wetlands, Hydrology

## Abstract

Temporary aquatic habitats are essential for maintaining regional fish diversity, especially in tropical ecosystems. However, artificial ephemeral environments such as roadside ditches remain largely overlooked in ecological research, despite their potential to support native, threatened and even invasive species. In this study, we first assess the fish community composition in roadside ditches of the Atlantic Forest, southeastern Brazil. These habitats exhibited extreme environmental conditions, including hypoxic waters, elevated temperature fluctuations and very acidic waters. Throughout the year, we recorded 17 fish species, including endangered and exotic taxa, across 36 ditches sampled in the Preto River microbasin. Species richness was higher during the Wet Period, likely due to increased hydrological connectivity with nearby streams. Although species richness was higher during the Wet Period, species composition did not differ between periods. Redundancy analysis revealed low explanation of environmental variables on community structure, while a significant positive correlation between spatial distance and community dissimilarity suggested dispersal limitation as a major structuring force. These findings highlight both the ecological relevance of artificial temporary habitats in fragmented landscapes and the requirement to integrate such environments into regional species inventories, especially as natural wetlands continue to decline.

## INTRODUCTION

Temporary aquatic habitats are dynamic ecosystems that often harbor high levels of endemism and a specialized ichthyofauna adapted to extreme environmental conditions (Williams, 2007). Survival in such ephemeral environments requires physiological and behavioral adaptations to cope with stressful environmental conditions, with several fish species exhibiting exceptional physiological tolerance to sharp fluctuations in water temperature and hypoxic (Polačik, Podrabsky, 2015), while others have evolved amphibious lifestyle to exploit these unstable habitats (Turko *et al*., 2021). In the Neotropical region, studies on temporary aquatic habitats have primarily focused on temporary pools formed by rainfall and river flooding. However, most research emphasizes taxonomic descriptions of new species (Costa, 2006; Drawert, 2022; Ramos *et al*., 2023), rather than understanding the structure and composition of fish communities, also an essential step for assessing ecological processes, evolutionary strategies, and conservation needs. Therefore, filling these knowledge shortfalls, particularly regarding species distribution and habitat associations, is critical (Freitas *et al*., 2021; Reis *et al*., 2024).

Despite their ecological importance, studies on fish community composition in temporary aquatic habitats remain scarce and are largely restricted to the Amazon region (Pazin *et al*., 2006; Espírito-Santo *et al*., 2009). In contrast, in the Atlantic Forest, research has predominantly focused on taxonomy, genetics, and population-level dynamics (Costa, 2002; 2006; 2009; Abilhoa *et al*., 2010; Contente, Stefanoni, 2010; Berbel-Filho *et al*., 2021; Guedes *et al*., 2023), with a notable absence of studies addressing community-level patterns in temporary environments. This lack of ecological knowledge is particularly concerning in a biome that is both a global biodiversity hotspot and home to approximately 36% of Brazil’s threatened small freshwater fish species (Castro, Polaz, 2020).

Among these temporary habitats, roadside ditches represent an overlooked but potentially important artificial temporary habitat, particularly in coastal restinga forests of the Atlantic Forest, where such temporary waters may host threatened or narrowly endemic species (Giongo *et al*., 2023; Costa *et al*., 2024a). Some fish species have even been documented exclusively in artificial habitats, such as highway gutters (Costa, 2002), reinforcing the urgent need to investigate these systems. Although artificial temporary environments are rarely addressed in fish community ecology, evidence from macroinvertebrate studies shows that even human-made ephemeral habitats, such as tire tracks, can harbor unique and rare species (Armitage *et al*., 2012). Similarly, a study conducted in Florida, USA, on fish assemblages in artificial ditch systems highlighted their potential for ecological research and conservation applications (Hohausová *et al*., 2010). Despite this, no study has yet examined fish community composition in artificial temporary habitats within the Neotropical region, leaving a significant gap in our understanding of fish living in these environments.

Given the rapid fragmentation of natural landscapes (Vancine *et al*., 2024), artificial ephemeral habitats, though often dismissed, can serve as natural laboratories for understanding the processes that structure species composition and how community assembly in new habitats in the absence of natural ones. So, documenting fish communities in such habitats is critical not only for biodiversity assessments, but also for informing conservation actions in a changing and increasingly fragmented world, since highly threatened species can use these artificial environments (Costa *et al*., 2024a), and also to investigate possible pathways of exotic species introductions (Brisson *et al*., 2010), since roadside ditches are human made and highly connected to human activity, favoring high tolerant invasive species that can cope with these extreme environments.

In this study we investigate the species composition of fish communities from roadside ditches in the Preto River microbasin, Itanhaém-SP, Brazil, an area dominated by blackwater streams flowing through restinga forests of the coastal Atlantic Forest. While the ichthyofauna of permanent streams in this basin, and adjacent areas, has been previously documented (Ferreira, Petrere, 2009; Ferreira *et al*., 2014; Giongo *et al*., 2023), the fish communities in temporary habitats remain entirely unexplored. Thus, we aimed to: (i) describe the fish species inhabiting these ephemeral artificial environments; (ii) assess differences in species richness and composition between Wet and Drier periods, given the seasonal influence of rainfall on hydrological connectivity; and (iii) examine how environmental conditions and spatial proximity between ditches influence community structure. We hypothesize that species richness and composition differ between rainfall periods, with higher richness during the Wet Period due to increased connectivity with nearby streams, potentially facilitating colonization by stream-dwelling species. In contrast, the Drier Period is expected to harbor mainly resident species adapted to extreme conditions, such as non-seasonal killifishes and air-breathing catfishes.

## METHODS

### Study area and sampling

The study was conducted in the Preto River microbasin, part of the Itanhaém River basin, in the coastal Atlantic Forest of São Paulo State, Brazil (Figure 1A). The Itanhaém River basin, the second largest coastal basin in the state, is situated in the central-southern region of São Paulo’s coastline. The Preto River microbasin lies predominantly within the coastal plain, characterized by sandy-clay soils and remnant restinga forests. Its waters are dark, with high levels of dissolved organic carbon (DOC) resulting from the microbial decomposition of lignin-rich materials. These blackwaters are also highly acidic and exhibit very low oxygen concentrations (Camargo, Cancian, 2016).

**Figure 1.**
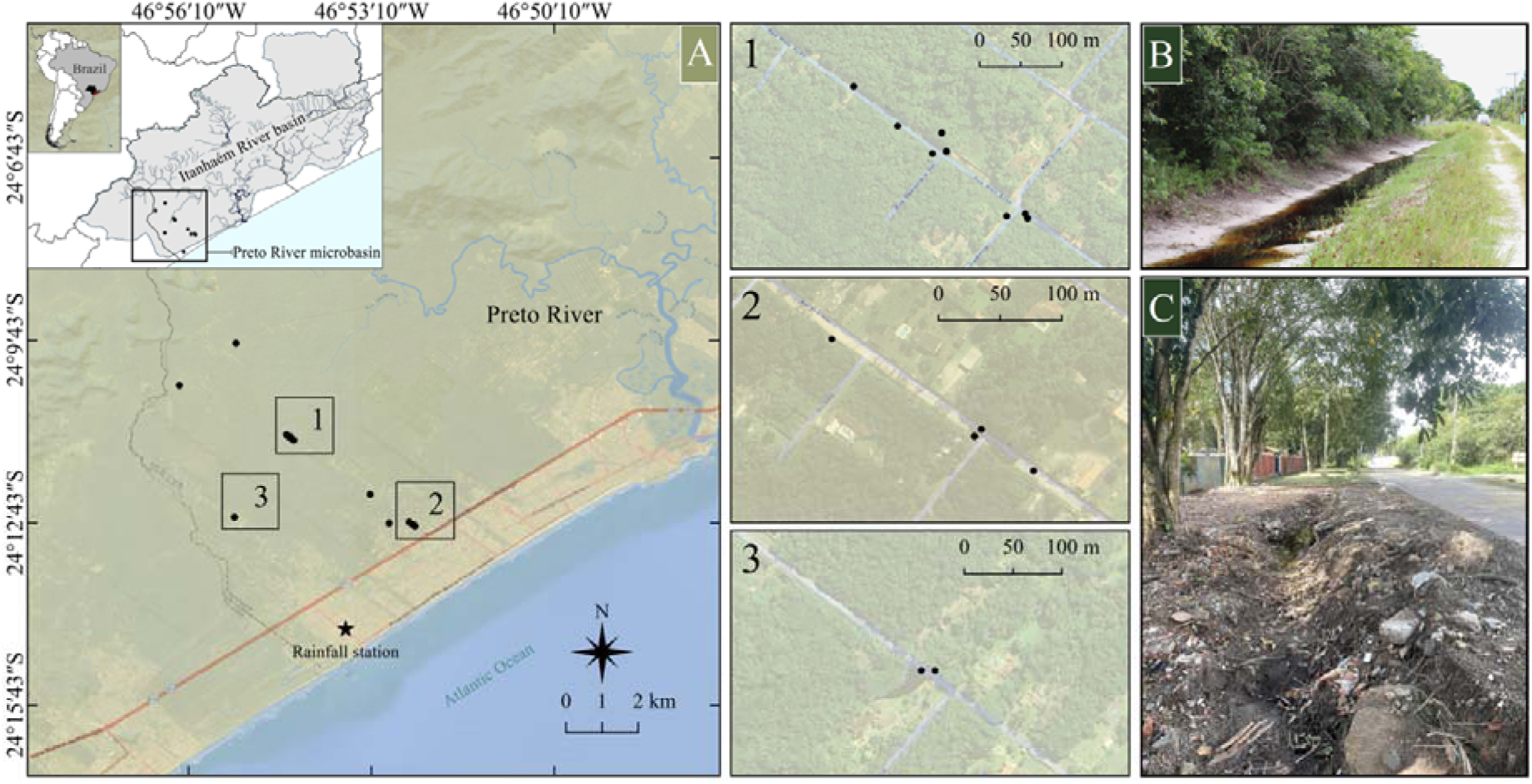
(A) Sampled roadside ditches at the Preto River microbasin, part of the Itanhaém River basin, Itanhaém-SP, Brazil. (B) Roadside ditch filled with water after stream flooding and rainfall. (C) Another roadside ditch dried after the water recedes.

The fish composition and distribution in the streams of the Itanhaém River basin have been documented by Ferreira, Petrere (2009), and Ferreira *et al*. (2014), who identified distinct fish zones within this important segment of the Atlantic Forest’s coastal basins, marked by high endemism. This endemism stems from the region’s natural isolation, with mountainous terrain acting as a physical barrier upstream and downstream by the Atlantic Ocean (Menezes *et al*., 2007; Giongo *et al*., 2023). However, no information is available regarding the fish fauna inhabiting temporary habitats in this region, particularly artificial temporary environments such as roadside ditches. These ditches are excavated alongside unpaved roads and have been maintained for several years (Figure 1B). They are temporarily flooded by rainfall and overflow from nearby streams, followed by a dry phase when water recedes (Figure 1C).

A total of 36 roadside ditches were sampled across the Preto River microbasin from January to December 2024, with three different ditches sampled each month. The same ditch (or another within the same temporarily flooded area) was only resampled in a later month if it had undergone complete drying followed by full reflooding, ensuring the site could be recolonized naturally. This criterion was adopted to avoid carryover effects from previous sampling on the structure of newly colonized communities in these artificial habitats. All sampling locations were georeferenced using GPS to allow continuous monitoring. Although 36 samples were obtained over the year, the monthly samples were deliberately limited to minimize disturbance in this ecosystem with highly vulnerable species. This precaution was reinforced after a Critically Endangered killifish species was rediscovered at one roadside ditch during the first three months of sampling (Costa *et al*., 2024a), with the joint occurrence of other species with a high level of threat (Costa *et al*., 2024b).

To standardize sampling effort across sites, each ditch was sampled for 15 minutes by two researchers using hand nets with an oval mouth (50 x 40 cm) with a 1 mm mesh size, appropriate for capturing small-bodied fish. The sampling method was specifically designed for this study, as traditional seines were impractical due to the abundance of submerged branches and human litter in these habitats. The chosen method is also well-suited for small aquatic environments, facilitating future comparisons with other temporary habitats, such as temporary pools. This is particularly relevant given the lack of previous studies investigating fish community composition in similar artificial environments in the Neotropical region.

Following sampling, specimens were anesthetized using 1 to 1.5 mL of eugenol per liter of water, fixed in 10% formalin for 48 hours, and subsequently preserved in 70% ethanol. All procedures were approved by the Ethics Committee for the Use of Animals at São Paulo State University (UNESP) – Bioscience Institute (CEUA – IB/CLP no 15/2023), and sampling was authorized under SISBIO license 90241-1. Collected specimens were identified and deposited in the Fish Collection (DZSJRP) of UNESP in São José do Rio Preto.

In the field, environmental variables were recorded at each site, including maximum length (m), maximum width (m), maximum depth (m), and distance to the nearest stream (m), with an open reel fiberglass measuring tape. Distance to the nearest stream was measured directly using a reel tape only when the stream was nearby, or obtained from Google Maps satellite imagery for distances greater than 100 meters. Physicochemical parameters were also recorded using a Hanna portable multiprobe (HI98194), including pH, temperature (°C), dissolved oxygen (mg.L^-1^), conductivity (µS.cmL¹), and total dissolved solids (mg.L^-1^). The water volume of each ditch (m³) was estimated assuming a half-ellipsoid shape, using the formula: *V* = 2/3 *x* π *x a x b x c*, where *V* is the volume, *a* is the maximum length (m), *b* is the maximum width (m), and *c* is the maximum depth (m).

Rainfall patterns, which play a crucial role in the dynamics of temporary environments, were used to classify months into two distinct periods: the Wet Period, encompassing November to March, and the Drier Period, from April to October. Although the dry season in Brazilian coastal streams is not strongly pronounced due to consistent rainfall throughout the year (Tonhasca, 2005), regional precipitation data indicate monthly variations in rainfall intensity, justifying this classification to better understand patterns in temporary environments, which remain largely unexplored. Therefore, we decided to use the term Drier Period instead of Dry Period because even in the period of less rainfall, there is still considerable rain in the region.

Rainfall data were obtained from the National Center for Monitoring and Alerts for Natural Disasters (CEMADEN), specifically from the Balneário Gaivota station (24°14’27.6”S, 46°53’34.8”W) in Itanhaém-SP, which is located near the sampled sites. The classification of each month into two rainfall periods was based on the mean monthly rainfall (mm) (Figure 2), using monthly precipitation data from 2016 to 2024, with months with an average greater than 150 mm being grouped in the Wet Period. To ensure data reliability, raw 10-minute interval rainfall records were aggregated into hourly sums. A day was excluded from analysis if it contained fewer than 18 hours of available functional rainfall data. Likewise, a month was excluded from the final monthly precipitation sum if fewer than 20 days met this data availability criterion.

**Figure 2.**
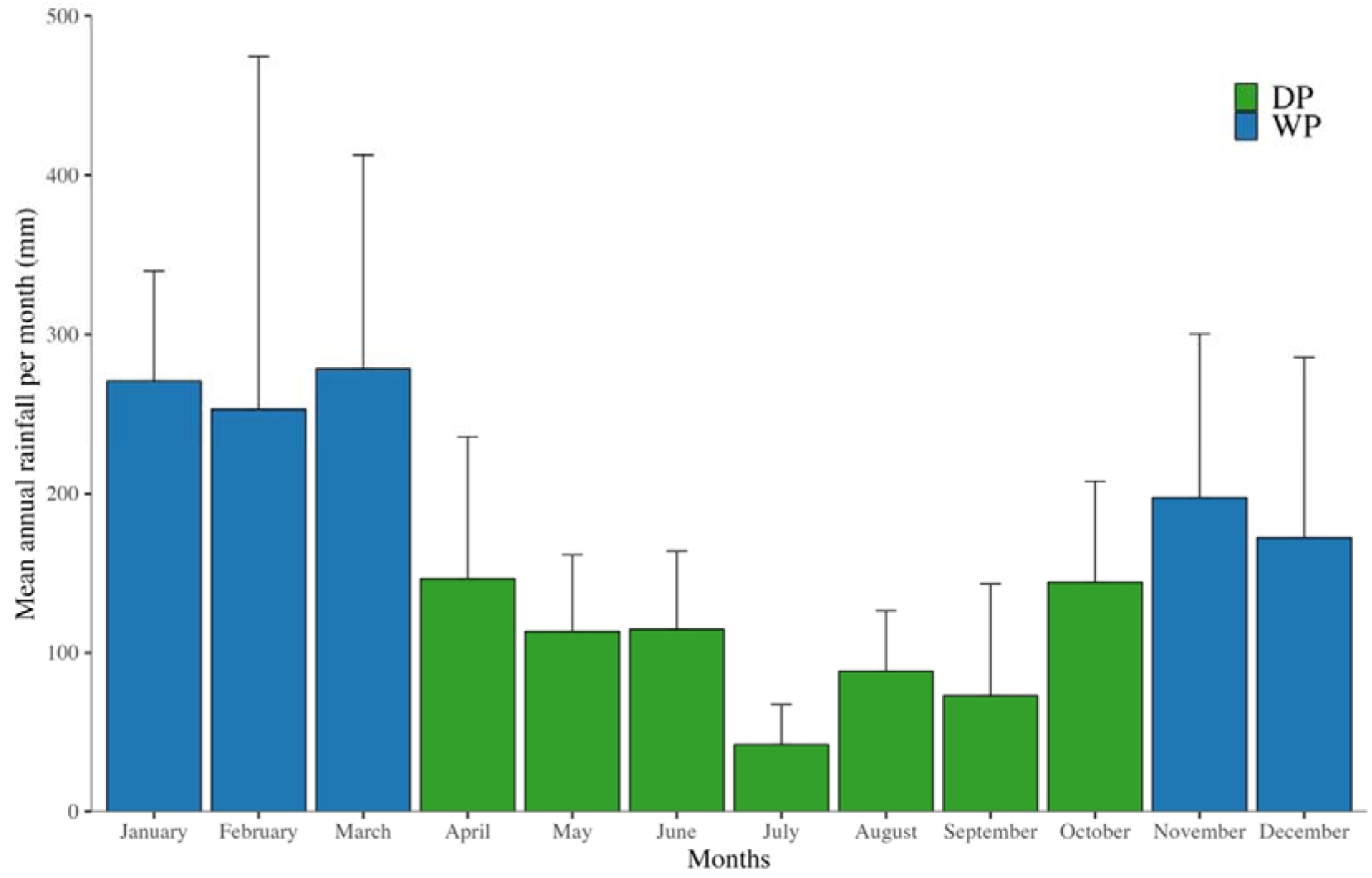
Mean annual rainfall per month (mm) in the Preto River microbasin region, Itanhaém-SP, Brazil, based on monthly precipitation data from 2016 to 2024. Bars represent the mean monthly rainfall, with error bars indicating standard deviations. Months were classified as part of the Drier Period (DP) or Wet Period (WP). Data were obtained from the National Center for Monitoring and Alerts for Natural Disasters (CEMADEN), using records from the Balneário Gaivota station (24°14’27.6”S, 46°53’34.8”W).

### Statistical analysis

To explore the relationships among environmental variables across the Drier and Wet periods a Principal Component Analysis (PCA) was conducted, including the following environmental variables: volume (m³), temperature (°C), pH, dissolved oxygen (mg.L^-1^), stream distance (m), total solids (mg.L^-1^) and conductivity (µS.cm^-1^). The variables maximum length (m), maximum width (m), and maximum depth (m) were excluded from the analysis because they were used to calculate the volume of the water bodies, avoiding redundancy.

Prior to the analysis, all environmental variables were standardized (mean = 0, standard deviation = 1) to ensure comparability among variables measured on different scales.

Also, to assess and compare the species richness between the periods rarefaction and extrapolation curves were generated using the *iNEXT* package in R (Hsieh *et al*., 2016). This analysis was based on incidence data, which accounts for species presence/absence across sampling units, and was performed using Hill numbers (q = 0), which corresponds to species richness. Rarefaction and extrapolation were estimated up to 40 sampling units to project the species accumulation. The variability in the estimates was assessed using 999 bootstrap replications, providing 95% confidence intervals.

We also performed a Permutational Multivariate Analysis of Variance (PERMANOVA) using the Bray-Curtis dissimilarity index to assess differences in fish community composition between the two periods, and the analysis was conducted with 999 permutations to evaluate the statistical significance. To verify the assumption of homogeneity of multivariate dispersions we applied a Permutational Multivariate Analysis of Dispersion (PERMDISP) using the Bray-Curtis distance matrix. This step ensured that any detected difference was not due to differences in group dispersion. To visualize patterns in fish community composition between the periods a Non-metric Multidimensional Scaling (NMDS) ordination was performed based on the Bray-Curtis dissimilarity index. The NMDS was computed using two dimensions (k = 2) to effectively represent the multivariate space, with a maximum of 100 random starts to ensure convergence to a global solution. To assess the fit of the ordination, the stress value was used as a measure of goodness-of-fit, with values below 0.2 considered acceptable for ecological data interpretation.

To explore how environmental variables influence fish community composition a Redundancy Analysis (RDA) was performed using the Bray-Curtis dissimilarity index. Prior to the analysis, the community matrix was transformed using the Hellinger transformation to minimize the impact of rare species and zero inflation, which is appropriate for abundance data. Environmental variables were selected based on the Variance Inflation Factor (VIF), with a threshold of VIF < 3 to avoid multicollinearity. The final model included the following variables: stream distance (m), volume (m³), dissolved oxygen (mg.L^-1^), pH and temperature (°C).

We also conducted a Mantel test to investigate the influence of spatial structure on fish community composition by assessing the correlation between geographic distances and community dissimilarity. Geographic coordinates from each sampling site were used to generate a pairwise geographic distance matrix (in kilometers). After that, the Hellinger transformed community matrix was used to calculate the Bray-Curtis dissimilarity to represent pairwise differences in community composition. The Mantel test was performed with 999 permutations to assess the significance of the correlation between geographic distance and community dissimilarity. PERMANOVA, PERMDISP, NMDS, RDA and Mantel test analyses were performed using the *vegan* R package (Oksanen *et al*., 2022). In addition, all raw community data, environmental variables, and coordinates of each sampling point, as well as code with the step-by-step data analysis, are available in a Git Hub repository for better reproducibility (https://github.com/JH-All/fish_community_roadside_ditches).

## RESULTS

The studied roadside ditches exhibited distinct environmental properties expected for such artificial habitats, including extremely low dissolved oxygen levels, ranging on average from 1.74 mg/L during the Wet Period (WP) to 0.34 mg/L in the Drier Period (DP). Additionally, extreme temperature oscillation was recorded throughout the year, reaching a maximum of 32.95°C (WP) and a minimum of 16.14°C (DP), along with substantial variability in water volume (on average 27.36 and 56.88 m^3^ in DP and WP, respectively). The pH ranged from very acidic (2.78) to circum-neutral (6.49) with average of 4.86 in WP and 5.20 in DP (Table 1). There was partial overlap in environmental properties between the periods; however, it was possible to observe higher values of dissolved oxygen, temperature, and volume during the Wet Period samples when compared to the Drier Period (Figure S1).

**Table 1.** Summary of environmental variables measured in roadside ditches during the Wet Period and Drier Period in the Atlantic Forest. Values represent minimum (Min), maximum (Max), mean and standard deviation (SD) for each variable (N=36).

| Variables | Wet Period |  |  | Drier Period |  |  |
| --- | --- | --- | --- | --- | --- | --- |
|  | Min | Max | Mean (SD) | Min | Max | Mean (SD) |
| Distance to nearest stream (m) | 0.30 | 851 | 206.14 (292.70) | 0.30 | 703.08 | 367.61 (313.07) |
| Maximum length (m) | 2.20 | 57.65 | 23.93 (15.61) | 1.12 | 49.10 | 20.25 (14.87) |
| Maximum width (m) | 0.81 | 9.0 | 2.75 (1.82) | 1.10 | 4.70 | 2.23 (0.80) |
| Maximum depth (m) | 0.11 | 0.80 | 0.28 (0.15) | 0.13 | 0.80 | 0.32 (0.17) |
| Volume (m<sup>3</sup>) | 1.49 | 293.40 | 56.88 (78.49) | 0.63 | 165.37 | 27.36 (34.94) |
| Temperature (°C) | 20.89 | 32.95 | 26.78 (3.11) | 16.14 | 27.06 | 21.47 (3) |
| Dissolved oxygen (mg.L<sup>-1</sup>) | 0 | 5.20 | 1.74 (1.75) | 0 | 2.62 | 0.34 (0.78) |
| pH | 2.78 | 6.30 | 4.86 (1.09) | 3.33 | 6.49 | 5.20 (1.03) |
| Conductivity ($\mu\text{S.cm}^{-1}$) | 47 | 162 | 85.13 (32.26) | 48.6 | 154 | 100.74 (40.17) |
| Total Dissolved Solids ($\text{mg.L}^{-1}$) | 30 | 105 | 54.20 (21.87) | 26 | 77 | 50.52 (17.91) |

A total of 787 individuals were collected, representing 17 species from 10 families and five orders (Table 2). The most abundant species was *Atlantirivulus santensis* (Köhler, 1906) (N = 212), followed by *Hyphessobrycon boulengeri* (Eigenmann, 1907) (N = 131), *Phalloceros reisi* Lucinda, 2008 (N = 104), *Poecilia reticulata* Peters, 1859 (N = 77), *Hyphessobrycon bifasciatus* Ellis, 1911 (N = 72), and *Mimagoniates lateralis* (Nichols, 1913) (N = 62). Several species were exclusively recorded during the Wet Period, including *Hollandichthys multifasciatus* (Eigenmann & Norris, 1900), *Geophagus iporangensis* Haseman, 1911, *Hoplosternum littorale* (Hancock, 1828), *Scleromystax macropterus* (Regan, 1913) and *Synbranchus marmoratus* Bloch, 1795. In contrast, only *Pseudotothyris obtusa* (Miranda Ribeiro, 1911) was exclusive to the Drier Period (Table 2).

**Table 2.** Fish species recorded in 36 roadside ditch samples in the Atlantic Forest during the Wet Period (WP) and Drier Period (DP). The table presents the number of individuals sampled in each period, the total abundance, and the corresponding voucher numbers deposited in the DZSJRP collection. Species marked with an asterisk (*) are classified as exotic. Threatened species are marked with their category in superscript. VU = Vulnerable. EN = Endangered. CR = Critically Endangered. Conservation status of each species was based on the Biodiversity Extinction Risk Assessment System – SALVE (ICMBio, 2025) and in state decrees (Estado de São Paulo, 2014).

| Order/Family | Species | WP | DP | Total | Voucher<br>DZSJRP |
| --- | --- | --- | --- | --- | --- |
| CHARACIFORMES |  |  |  |  |  |
| Stevardiidae |  |  |  |  |  |
|  | *Mimagoniates lateralis* (Nichols, 1913)<sup>VU</sup> | 41 | 21 | 62 | 25059 |
| Acestorhamphidae |  |  |  |  |  |
|  | *Hyphessobrycon boulengeri* (Eigenmann, 1907) | 45 | 86 | 131 | 25058 |
|  | *Hyphessobrycon bifasciatus* Ellis, 1911 | 18 | 54 | 72 | 25070 |
|  | *Rachoviscus crassiceps* Myers, 1926 <sup>CR</sup> | 1 | 8 | 9 | 23340 |
|  | *Hollandichthys multifasciatus* (Eigenmann & Norris, 1900) | 1 | - | 1 | 25062 |
| Erythrinidae |  |  |  |  |  |
|  | *Hoplias malabaricus* (Bloch, 1794) | 5 | 3 | 8 | 25065 |
| Lebiasinidae |  |  |  |  |  |
|  | *Nannostomus beckfordi* Günther, 1872 <sup>*</sup> | 29 | 6 | 35 | 25069 |
| CICHLIFORMES |  |  |  |  |  |
| Cichlidae |  |  |  |  |  |
|  | *Geophagus iporangensis* Haseman, 1911 | 4 | - | 4 | 25066 |
| CYPRINODONTIFORMES |  |  |  |  |  |
| Poeciliidae |  |  |  |  |  |
|  | *Phalloceros reisi* Lucinda, 2008 | 42 | 62 | 104 | 25064 |
|  | *Poecilia reticulata* Peters, 1859 <sup>*</sup> | 4 | 73 | 77 | 25068 |
| Rivulidae |  |  |  |  |  |
|  | *Atlantirivulus santensis* (Köhler, 1906) | 86 | 126 | 212 | 25056 |
|  | *Leptopanchax itanhaensis* (Costa, 2008) <sup>CR</sup> | 8 | 5 | 13 | 24845; 24846 |
| SILURIFORMES |  |  |  |  |  |
| Callichthyidae |  |  |  |  |  |
|  | *Callichthys callichthys* (Linnaeus, 1758) | 45 | 4 | 49 | 25057 |
|  | *Hoplosternum littorale* (Hancock, 1828) | 1 | - | 1 | 25173 |
|  | *Scleromystax macropterus* (Regan, 1913) <sup>EN</sup> | 1 | - | 1 | 25060 |
| Loricariidae |  |  |  |  |  |
|  | *Pseudotothyris obtusa* (Miranda Ribeiro, 1911) | - | 6 | 6 | 25073 |
| SYNBRANCHIFORMES |  |  |  |  |  |
| Synbranchidae |  |  |  |  |  |
|  | *Synbranchus marmoratus* Bloch, 1795 | 2 | - | 2 | 25072 |

Two exotic species appeared with notable abundance: *Nannostomus beckfordi* Günther, 1872, recorded for the first time in temporary habitats of the region, and *P. reticulata*, marking its first record in the basin. Although the majority of the native species are classified as Least Concern (LC), one is categorized as Vulnerable (VU) – *M. lateralis*, one as Endangered (EN) – *S. macropterus*, and two as Critically Endangered (CR) – *Leptopanchax itanhaensis* (Costa, 2008) and *Rachoviscus crassiceps* Myers, 1926 (Table 2). Photographs of all species are provided in Figure 3.

**Figure 3.**
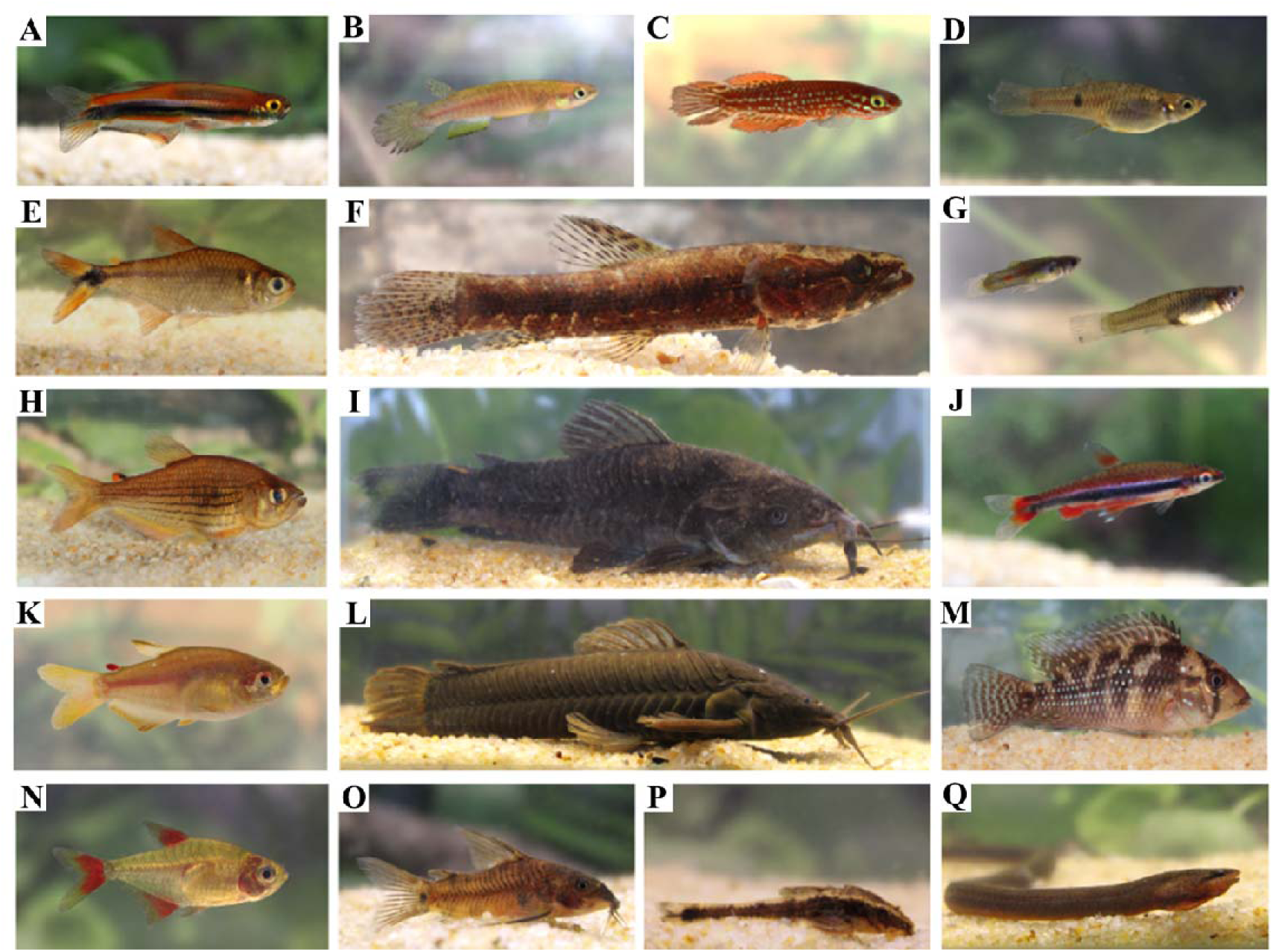
Species sampled in the roadside ditches from the Preto River microbasin, Itanhaém-SP. (A) *Mimagoniates lateralis*. (B) *Atlantirivulus santensis*. (C) *Leptopanchax itanhaensis*. (D) *Phalloceros reisi*. (E) *Hyphessobrycon boulengeri*. (F) *Hoplias malabaricus*. (G) *Poecilia reticulata*. (H) *Hollandichthys multifasciatus*. (I) *Hoplosternum littorale*. (J) *Nannostomus beckfordi*. (K) *Rachoviscus crassiceps*. (L) *Callichthys callichthys*. (M) *Geophagus iporangensis*. (N) *Hyphessobrycon bifasciatus*. (O) *Scleromystax macropterus*. (P) *Pseudotothyris obtusa*. (Q) *Synbranchus marmoratus*. Photographs of live specimens taken by Amanda Selinger during fieldwork.

Species richness increased more rapidly during the Wet Period compared to the Drier Period as sampling effort expanded. While species accumulation in the Drier Period showed signs of stabilization close to 10 interpolated samples, the Wet Period continued to show potential for additional species even after 15 samples (Figure 4). Despite these patterns, the community composition did not differ significantly between periods (PERMANOVA: F = 1.08, R² = 0.03, p = 0.34; PERMDISP: F = 0.06 p = 0.80). This result is further supported by the NMDS, which revealed a partial overlap in species composition between the periods (Figure 5A). However, some individual sampling points appeared to differ, with a slight association of *C. callichthys*, *L. itanhaensis*, and *R. crassiceps* with the Wet Period (Figure 5B).

**Figure 4.**
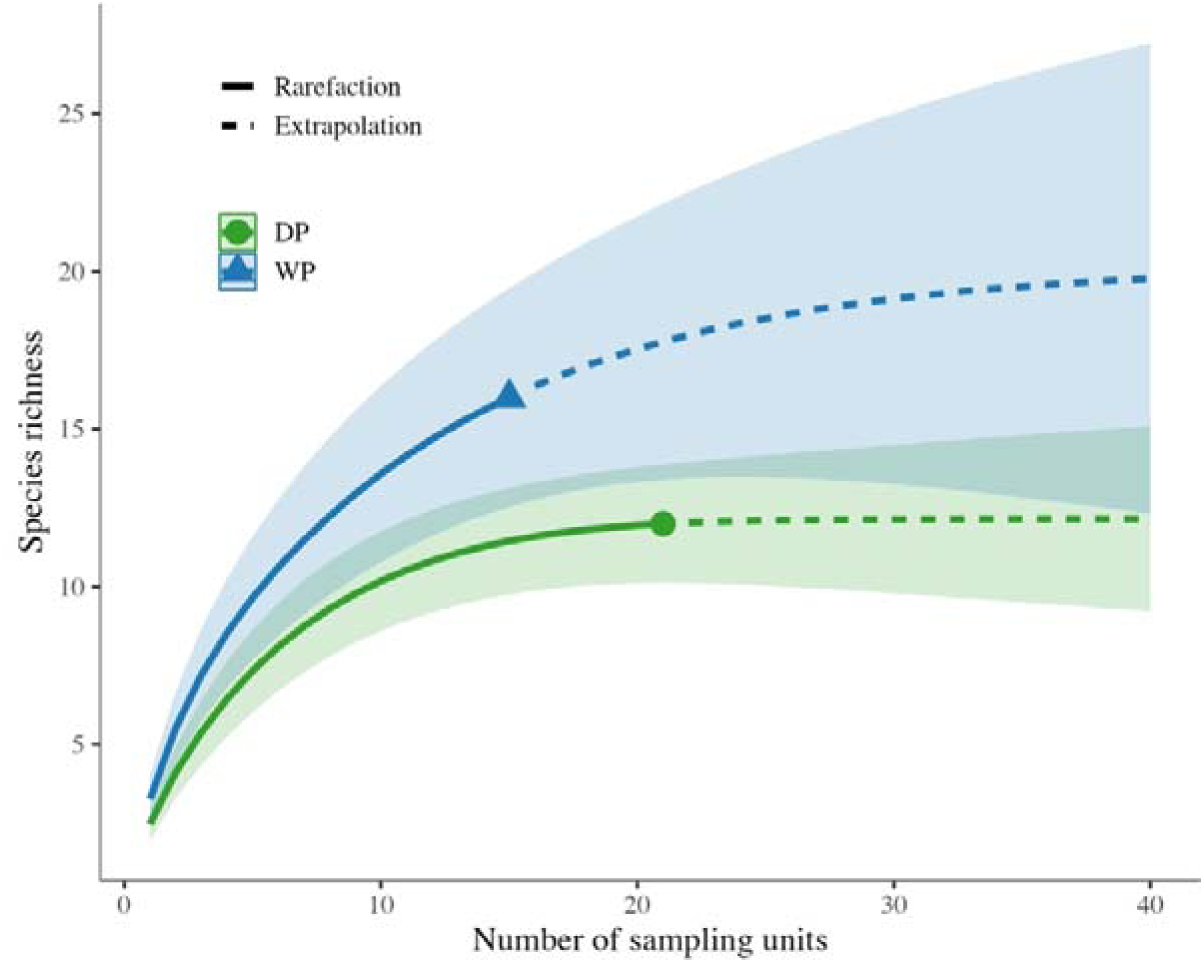
Rarefaction and extrapolation curves of fish species richness in roadside ditches during the Wet Period (WP) and Drier Period (DP) in the Atlantic Forest. Solid lines represent the rarefaction curves based on the observed data, while dashed lines indicate the extrapolated species richness beyond the observed sampling effort. Shaded areas correspond to the 95% confidence intervals.

**Figure 5.**
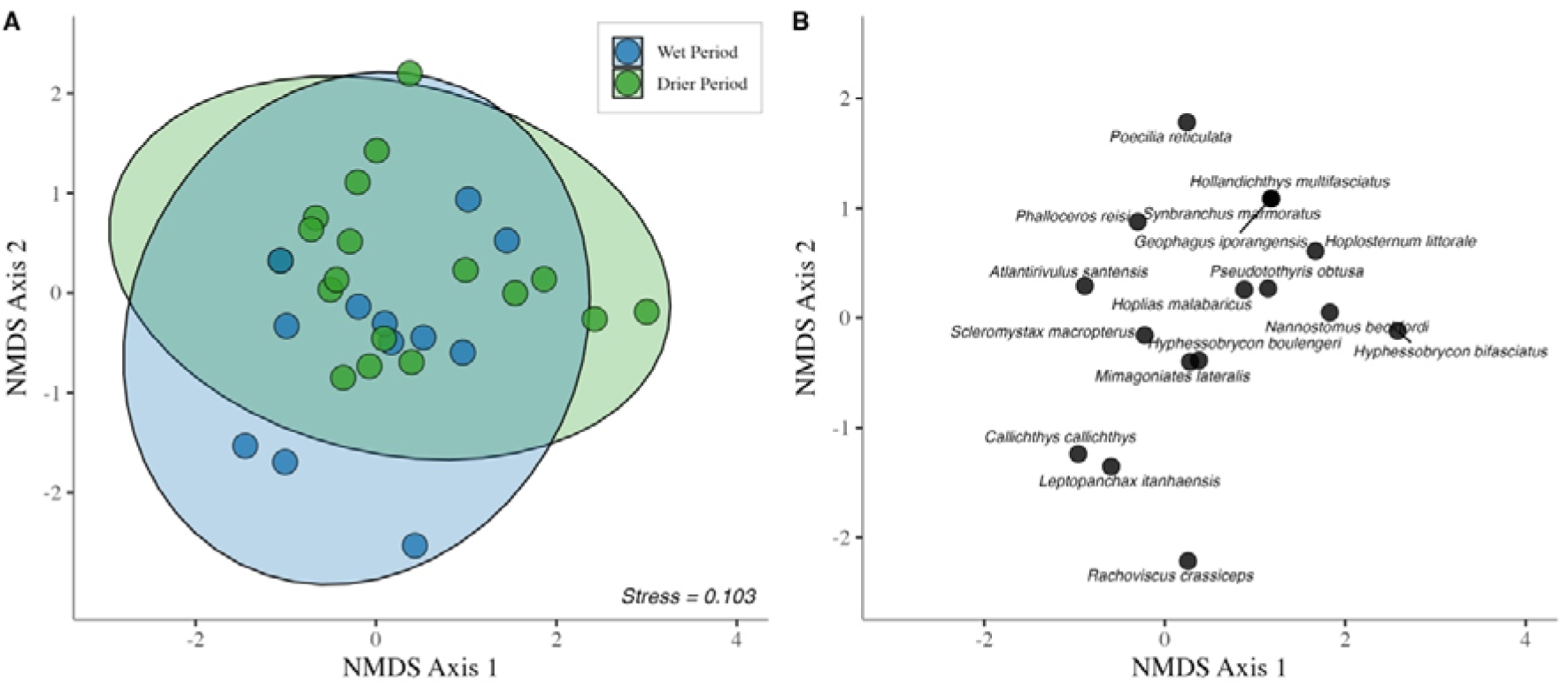
Non-metric Multidimensional Scaling (NMDS) ordination of fish assemblages in roadside ditches during the Wet Period and Drier Period in the Atlantic Forest. (A) Distribution of sampling sites for both periods, where overlapping points show greater similarity. Ellipses represent 95% confidence intervals for each period. (B) Positioning of fish species within the ordination space.

The RDA revealed that environmental variables collectively explained only a modest portion of the variation in fish community composition, with RDA1 accounting for 14.44% and RDA2 for 5.31% of the total variance. Despite the relatively low explained variance, the observed patterns hold important ecological significance. For instance, increases in distance from the nearest stream were positively associated with the occurrence of *P. reticulata*, but showed a negative association with species such as *M. lateralis* and *H. boulengeri*. *Atlantirivulus santensis* exhibited a negative relationship with increasing water volume and a positive association with higher pH levels. Additionally, *N. beckfordi* and *M. lateralis* exhibited a small – but positive – association with higher levels of dissolved oxygen (Figure 6). However, a significant positive correlation was found between spatial distance among samples and community dissimilarity (Mantel r = 0.31, p = 0.001), suggesting that geographically closer sites tend to have more similar fish communities (Figure S2), where limited dispersion might exert an important role in structuring the community of these artificial temporary environments.

**Figure 6.**
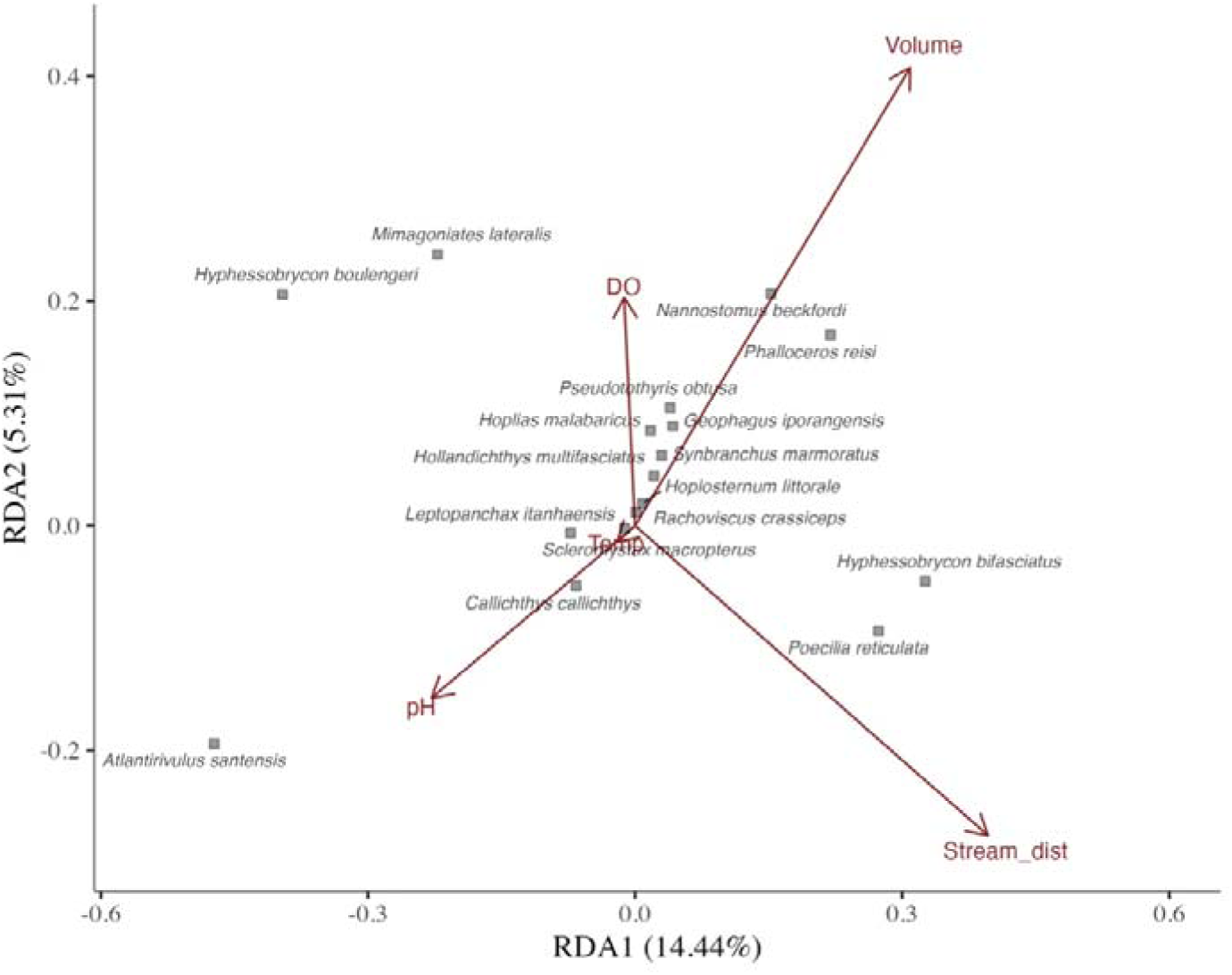
Redundancy Analysis (RDA) biplot showing the relationship between fish species composition and environmental variables in roadside ditches of the Atlantic Forest. Temp = Temperature (°C). DO = Dissolved oxygen (mg.L^-1^). Volume = Volume in m^3^. Stream_dist = Distance to nearest stream (m).

## DISCUSSION

In this study we provide the first description of the fish communities inhabiting roadside ditches in the Atlantic Forest, artificial and temporary habitats that, despite their instability, support a distinct and surprisingly diverse ichthyofauna. Even under extreme oscillation in the physicochemical conditions, as seen by dissolved oxygen and pH, and elevated temperatures during some periods of the year, we recorded a total of 17 fish species, including threatened taxa, and species likely introduced through human activities. Our results support the hypothesis that species richness increases during the Wet Period, likely due to enhanced hydrological connectivity with nearby streams. However, contrary to our expectations, species composition did not differ significantly between rainfall periods, regardless of the total richness observed. Finally, we also take initial steps toward understanding how environmental variables and spatial proximity between ditches shape fish communities in these overlooked ecologically unique environments.

The roadside ditches demonstrated harsh environmental conditions typical of extreme temporary habitats, particularly in terms of dissolved oxygen, which remained critically low throughout the year, averaging 0.32 mg/L during the Drier Period and 1.74 mg/L during the Wet Period. These values are substantially lower than those reported for blackwater streams in the region, which average 5.55 mg/L (Giongo *et al*., 2023), indicating a significant physiological challenge for fish dispersing from streams into ditches. Temperature oscillations were also marked, ranging from 16.14°C (DP) to 32.95°C (WP), far exceeding the thermal variability observed in nearby streams, ranging from 17°C to 22°C (Ferreira *et al*., 2014; Giongo *et al*., 2023). Notably, the maximum temperature recorded in ditches was only ∼5°C below thresholds linked to mass fish mortality events in the Amazon (Braz-Mota, Val, 2024), suggesting that extreme summer heatwaves – as seen during 2024 – might push many fish species close to their thermal limits. Although environmental conditions showed an overlap between periods (as indicated by the PCA), the Wet Period consistently exhibited on average higher oxygen levels, temperatures, and water volumes, likely reflecting the combined effects of increased rainfall and seasonal warming.

We recorded a surprisingly rich ichthyofauna for such a spatially restricted, ephemeral, and environmentally harsh habitat, with 17 fish species found in the roadside ditches. Of the 19 species previously documented in blackwater streams of the Preto River microbasin (Ferreira *et al*., 2014), 11 were also found in ditches, and more two other species that were reported in a stream fish zone that overlaps with the sampled area (*H. littorale*, *N. beckfordi*) (Ferreira, Petrere, 2009). Four additional species were recorded in ditches but had not been previously reported in community composition published in literature for this microbasin, detailed for the first time in this context (*L. itanhaensis*, *R. crassiceps*, *P. reticulata*, *H. bifasciatus*). These findings highlight the potential of artificial temporary habitats to contribute to regional species inventories, revealing both floodplain-associated taxa, highly threatened species and even introduced ones.

The most abundant species was *A. santensis*, a non-seasonal killifish broadly distributed along the São Paulo coast (Contente, Stefanoni, 2010). Although rarely abundant in stream communities (Ferreira, Petrere, 2009; Ferreira *et al*., 2014; Giongo *et al*., 2023), its high abundance in ditches suggests a preference for small flooded areas. The second and third most abundant species, *H. boulengeri* and *P. reisi*, respectively, are common in local streams and likely reach ditches during flooding, becoming trapped when water levels recede. Other stream-dwelling species, *H. multifasciatus*, *G. iporangensis*, *H. littorale*, *S. macropterus* and *S. marmoratus*, were observed only during the Wet Period, indicating that increased rainfall and flooding facilitate their entry into temporary habitats. This same aspect probably reflects the difference in accumulated species richness between periods, with the Wet Period presenting greater richness and delayed stabilization, when compared to the Drier Period, supporting our hypotheses.

However, remains unclear if this pattern reflects accidental drift or intentional use of flooded areas for reproduction, although previous studies suggests that some species may exploit these environments for reproduction (Espírito-Santo *et al*., 2013). Interestingly, *P. obtusa* was found only during the Drier Period. As a stream-associated species, its presence may reflect delayed movement or failure to detect environmental cues to exit temporary habitats, as observed in Amazonian species to avoid entrapment (Espírito-Santo *et al*., 2017). Such responses to hydroperiod and environmental variability can influence local species richness (Pazin *et al*., 2006) and likely play a role in shaping these ditch communities as well.

Although species richness varied between rainfall periods, our results did not support the hypothesis that species composition in roadside ditches differs significantly between the Wet and Drier periods, a pattern also reported in coastal streams (Tonhasca, 2005). However, the NMDS revealed non-random associations of certain species with the Wet Period, which, while not driving overall community turnover, suggest ecologically meaningful patterns. For instance, *L. itanhaensis*, an annual killifish recently rediscovered in this region (Costa *et al*., 2024a), was mostly found during the Wet Period. Although five individuals were recorded during the Drier Period, this may reflect the species life cycle, as even within the dry season, occasional rainfall events may trigger hatching.

A similar association was observed for *R. crassiceps*, a small tetra with a highly fragmented distribution and poorly understood ecology (Costa *et al*., 2024b), despite this, its total abundance was greater in the Drier Period, with the association found in the NMDS having to be interpreted with caution, since the species occurs in few sampling points, due to the very nature of its rarity. Despite its absence in regional stream surveys (Ferreira *et al*., 2014), it appears to favor flooded environments such as roadside ditches and temporary pools (Costa *et al*., 2024b), highlighting the need for further research on its habitat use, life history and persistence in ephemeral systems. *Callichthys callichthys*, another species associated with the Wet Period, possesses physiological traits such as aerial respiration – using intestine as air breathing organ – and terrestrial locomotion ability (Le Bail *et al*., 2000), which likely facilitate its dispersal across flooded landscapes during heavy rains.

The presence of threatened and endangered species in these artificial temporary habitats underscores the conservation relevance of considering these roadside ditches in species inventories. Species such as *L. itanhaensis* and *R. crassiceps* (Critically Endangered), *S. macropterus* (Endangered) and *M. lateralis* (Vulnerable) were recorded, suggesting that these environments may act as refugees or transient habitats. This reinforces the need for inventorying and monitoring artificial water bodies, especially before road paving, drainage works or routine maintenance, as such activities may inadvertently eliminate important species. At the same time, the occurrence of these species in artificial habitats raises concerns. It may reflect the loss or degradation of natural temporary environments, such as temporary pools in restinga forests, forcing species to occupy suboptimal or exposed habitats. These environments have extreme physicochemical conditions, and exotic species, as seen in our results, adding further stress to already vulnerable taxa. Future research should focus on assessing the impacts of pollution, biological invasions, and the adaptive physiological or behavioral mechanisms that enable these species to thrive in such challenging habitats.

The detection of the exotics *N. beckfordi* and *P. reticulata*, with the last one recorded for the first time in the entire Itanhaém River basin, raises additional conservation concerns. Notably, *P. reticulata* emerged as the fourth most abundant species in the community, highlighting its apparent tolerance to habitat degradation and pollution (Kennard *et al*., 2005; Silva *et al*., 2023). The RDA results further revealed a positive association between *P. reticulata* and greater distances from the nearest stream, suggesting that its presence is more frequent in more isolated roadside ditches, that are closer to areas with higher urbanization, where the occurrence of native species is likely lower. This spatial pattern may reflect intentional introductions, either by local residents or public authorities, as a mosquito control strategy. Such introductions are particularly worrisome, as subsequent flooding events could facilitate the dispersal of *P. reticulata* into nearby temporary pools or streams, potentially disrupting native fish communities through competition or disease transmission.

Although the RDA explained a limited portion of the variation in community composition, some ecologically meaningful patterns emerged. *Atlantirivulus santensis* showed a negative association with ditch volume and a positive association with pH, a pattern that may reflect the characteristics of smaller ditches filled by recent small rainfall events, which are less influenced by acidic floodwaters from nearby blackwater streams. The use of shallow, low-volume habitats by *A. santensis* aligns with its non-annual life cycle and potential for terrestrial jumps to escape predators and competition, as observed in other rivulid species (Espírito-Santo *et al*., 2019). Its amphibious lifestyle may also confer an advantage in hypoxic, ephemeral environments prone to desiccation (Turko *et al*., 2021). However, these behavioral and physiological traits still require further investigation in the context of this species and region.

Other species, such as *M. lateralis* and *H. boulengeri*, exhibited a negative relationship with distance from the nearest stream and a positive association with higher dissolved oxygen levels, consistent with their ecology as typical stream dwellers in the region (Ferreira *et al*., 2014), and with their physiology as pelagic species with high swimming activity level that requires a higher metabolic rate to sustain its lifestyle (Campos *et al*., 2018). Their presence in roadside ditches likely reflects dispersal during flooding, especially near permanent water bodies. Although some of these stream dwellers species are acidophilic (e.g., *M. lateralis* and *S. macropterus*), and thus potentially sensitive to the higher pH observed in some ditches, our data suggest that this factor does not act as a short-term environmental filter. Nonetheless, the effects of such novel conditions on their fitness remain an important question for future research.

Although environmental filtering often plays a key role in structuring fish assemblages in coastal streams (Silva *et al*., 2023), our results suggest a different dynamic in roadside ditches. The low explanatory power of environmental variables in the redundancy analysis, combined with a significant positive correlation between geographic distance and communities dissimilarity, indicates that dispersal limitation may be a dominant force shaping fish communities in these ephemeral habitats. For species lacking amphibious adaptations, colonization likely depends on complete flooding events, making geographically closer ditches more similar in species composition. This spatial structuring pattern resembles those observed in odonate communities in artificial ponds (McCauley, 2006) and in freshwater fish assemblages in lakes and reservoirs, where dispersal limitation has a greater influence on species richness than environmental heterogeneity (Drakou *et al*., 2009). While our findings represent an initial and small first step toward understanding fish metacommunity theory in artificial temporary habitats, further research on the topic is urgently needed. Studies focusing on both taxonomic and functional diversity, along environmental and spatial gradients, such as distance from streams, hydroperiod and temperature variation, are essential to clarify the mechanisms driving community assembly in these increasingly common and ecologically relevant systems.

Altogether, our findings offer a foundational understanding of fish community composition in artificial temporary habitats within a blackwater basin from the Atlantic Forest, revealing that even human-made environments with extreme physicochemical conditions can harbor high species richness, including taxa of conservation concern. This study represents the first step toward understanding how fish use such habitats in the absence of natural environments like temporary pools. As these artificial systems may be acting as refugees or ecological traps, their role in regional biodiversity dynamics deserves urgent attention. In a world facing accelerating habitat fragmentation, climate-driven hydrological shifts and widespread loss of natural wetlands, exploring how fish communities persist and adapt in ephemeral human-modified landscapes is not only timely but essential for guiding effective conservation strategies.

## ACKNOWLEDGMENTS

This study was financed, in part, by the São Paulo Research Foundation (FAPESP), Brasil. Process Number #2023/14344-5 and Coordenação de Aperfeiçoamento de Pessoal de Nível Superior - Brasil (CAPES) - Finance Code 001. We also thank the Núcleo de Pesquisas Hidrodinâmicas at Santa Cecília University team for the support with rainfall data. To the INCT-ADAPTA II project that is supported by CAPES (Finance Code 001), CNPq (#465540/2014-7) and FAPEAM (#06201187/2017). We would also like to thank Thomas Alves Vidal and Esli Mosna who provided support during the fieldwork.

