## Supplemental Figures for "Life on the road: fish communities composition in roadside ditches of the Atlantic Forest"

**SUPPLEMENTARY MATERIAL**

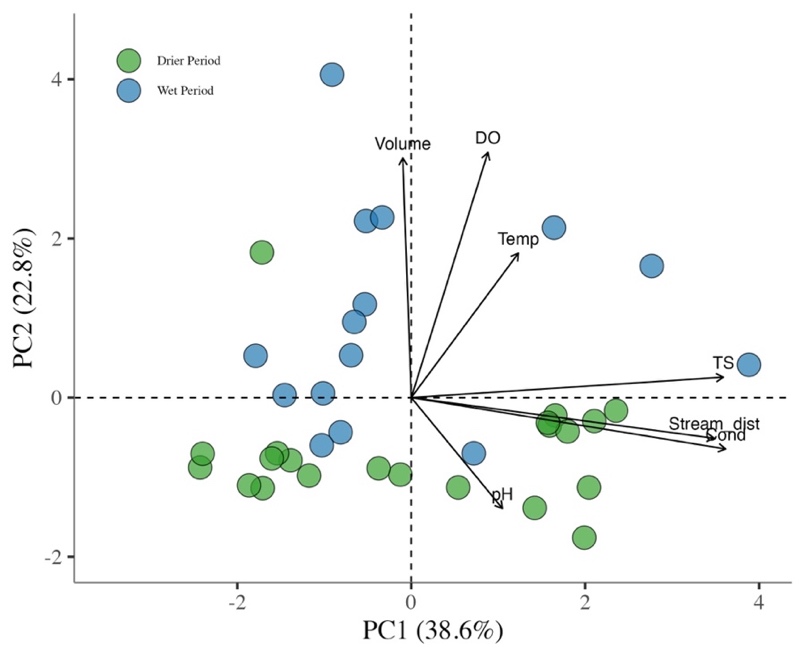

**Figure S1.** Principal Component Analysis (PCA) biplot illustrating the distribution of environmental variables in roadside ditches during the Wet Period (WP) (blue) and Drier Period (DP) (green). Temp = Temperature (°C). DO = Dissolved oxygen (mg.L^-1^). Volume = Volume in m^3^. Stream_dist = Distance to nearest stream (m). TS = Total Dissolved Solids (mg.L^-1^). Cond = Conductivity (μS.cm-^1^).

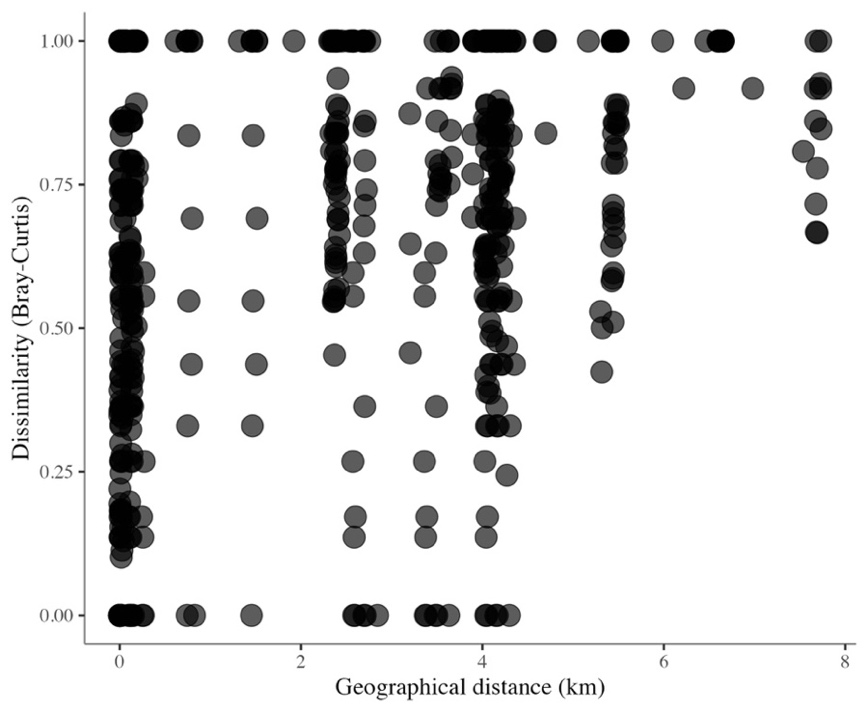

**Figure S2.** Relationship between geographic distance (in kilometers) and Bray-Curtis dissimilarity of fish communities in roadside ditches of the Atlantic Forest. Each point represents a pairwise comparison between two sites.
